# PlantAI: A Multi-Agent System for Plant Functional Genomics Analysis and Biological Knowledge Interpretation

**DOI:** 10.64898/2026.08.14.744760

**Authors:** Tingxun Wu, Zhuang Yang, Jin Shi, Meiling Zou, Yingdong Wu, Sirong Jiang, Chengcai Xia, Lin Kong, Lin Yang, Zhiqiang Xia

## Abstract

Plant functional genomics requires the integration of sequence, expression, evolutionary, regulatory and literature evidence. However, the corresponding analyses are often distributed across disparate programs, scripts and databases, creating substantial barriers to task organization and result interpretation. Here, we present PlantAI, a multi-agent system that integrates bioinformatics analysis, project-level process tracking and knowledge-assisted interpretation. A Main Agent coordinates two complementary routes: an analysis route that invokes bioinformatics tools for RNA-seq and gene-family analyses, and a knowledge route that uses PlantAI-RAG for knowledge retrieval and evidence synthesis. PlantAI-RAG currently contains 31,207 plant-science literature records, comprising approximately 3.82 million normalized entities and 8.25 million literature-supported relation assertions. In an evaluation using plant-science questions, it achieved a Gold evidence-assertion recall of 86.7%, while strict accuracy ranged from 77% to 82% across three independent evaluator models. We further demonstrate an end-to-end task using 24 rice RNA-seq libraries collected under salt stress, spanning transcriptome analysis, candidate-family screening, HXK/HKL family analysis and knowledge-assisted interpretation, and prioritize OsHXK8 for experimental validation. By preserving analysis artifacts, run manifests, logs and environment records, PlantAI supports result verification and repeat execution while linking project-derived results to traceable literature evidence. Together, these capabilities provide an integrated and auditable framework to support plant functional genomics research.

## 1. Introduction

Plant functional genomics seeks to connect evidence across sequence, expression, regulation, evolution and phenotype to identify genes, gene families and biological processes worthy of further validation. As plant genomic, transcriptomic and other omics data continue to accumulate, research increasingly extends beyond a single isolated statistical analysis. Expression changes usually need to be organized into candidate genes or candidate families; these candidates must then be placed in the contexts of phylogeny, conserved domains, gene structure, synteny, regulatory elements and protein interactions; and the resulting computational outputs must be compared with existing literature and database knowledge. A differential-expression table, a phylogenetic tree or a single knowledge question is therefore usually only one component of a complete research chain. Plant knowledge is itself dispersed across a growing body of papers, databases and unstructured text, increasing the difficulty of cross-source integration and evidence tracking ^1–4^.

Bioinformatics analysis has become an important component of plant-science research, but substantial barriers remain in practice. Many established tools depend on command-line environments and use different programming languages, input formats, software dependencies and parameter systems. Researchers must not only select appropriate programs but also configure environments, convert formats, coordinate stages, monitor run status and organize results. By integrating common analysis and visualization functions, TBtools-II showed that lowering barriers to tool use can directly support plant researchers working with large-scale biological data ^5^. However, when a research question spans RNA-seq, candidate screening, gene-family characterization and literature interpretation, integrating individual tools does not fully address task planning, cross-stage data transfer, project-result tracking and evidence organization. Multi-agent systems have recently been applied to bioinformatics task decomposition, workflow organization and specialized knowledge invocation ^6,7^. Specialist agents such as GeneAgent have also demonstrated the feasibility of combining domain databases with gene-set analysis and result self-verification ^8^. Plant-domain question-answering systems, meanwhile, have begun to use domain corpora, knowledge graphs and retrieval-augmented generation to improve the specialization and traceability of responses ^1–3^. Nevertheless, analysis-execution and knowledge-question-answering systems often remain separate: the former generates project results, whereas the latter answers general knowledge questions, with no continuous mechanism for bringing actual analysis artifacts back into literature retrieval and biological interpretation.

Against this background, we developed PlantAI, a multi-agent system for plant functional genomics research. PlantAI uses the Main Agent as its interaction and decision centre, routing user research tasks to either the Analysis Route or the Knowledge Route. Along the Analysis Route, the Intent Agent, Planning Agent and Execution Agent identify the analytical intent, construct an inspectable stage plan and dispatch confirmed analysis tasks, respectively. Researchers can confirm or revise plans and parameters, while the underlying computations are performed by a Bioinformatics Tool Layer comprising command-line modules, analysis scripts and external software. Along the Knowledge Route, the Knowledge Agent (PlantAI-RAG) decomposes questions, plans retrieval and synthesizes evidence, invokes the Knowledge Retrieval Layer for vector and knowledge-graph retrieval, and returns source-linked evidence to the Main Agent. Analysis results are stored in a Project workspace, enabling the Main Agent to read existing artifacts and combine them with literature evidence returned by PlantAI-RAG when composing project-aware biological interpretations. This paper presents the PlantAI architecture and its evidence-grounded Knowledge Agent, together with an end-to-end case that proceeds from rice salt-stress RNA-seq analysis through candidate-family screening and HXK/HKL family characterization to RAG-assisted interpretation. The case demonstrates how PlantAI organizes established bioinformatics methods, reproducible project records and knowledge retrieval into a continuous research workflow. Using rice salt stress as an example, it illustrates how PlantAI can connect candidate-gene screening, biological interpretation and subsequent experimental research in plant functional genomics while organizing traceable computational outputs and literature evidence.

## 2. Results

### 2.1 PlantAI multi-agent system and reproducible execution

PlantAI accepts research questions, knowledge questions and omics data, and the Main Agent determines whether each request should enter the Analysis Route or the Knowledge Route (Fig. 1). Along the Analysis Route, the Intent Agent identifies the analysis objective, the Planning Agent constructs a plan from project files and stage dependencies, the researcher may confirm or revise parameters and biological scope, and the Execution Agent passes the confirmed task to the Bioinformatics Tool Layer. This layer comprises CLI modules, analysis scripts and external software and performs the underlying computations, including quality control, alignment, quantification, statistical testing, phylogenetic inference and network construction. Large language models are used to understand, plan and organize tasks, but do not replace these analysis programs.

**Figure 1.**
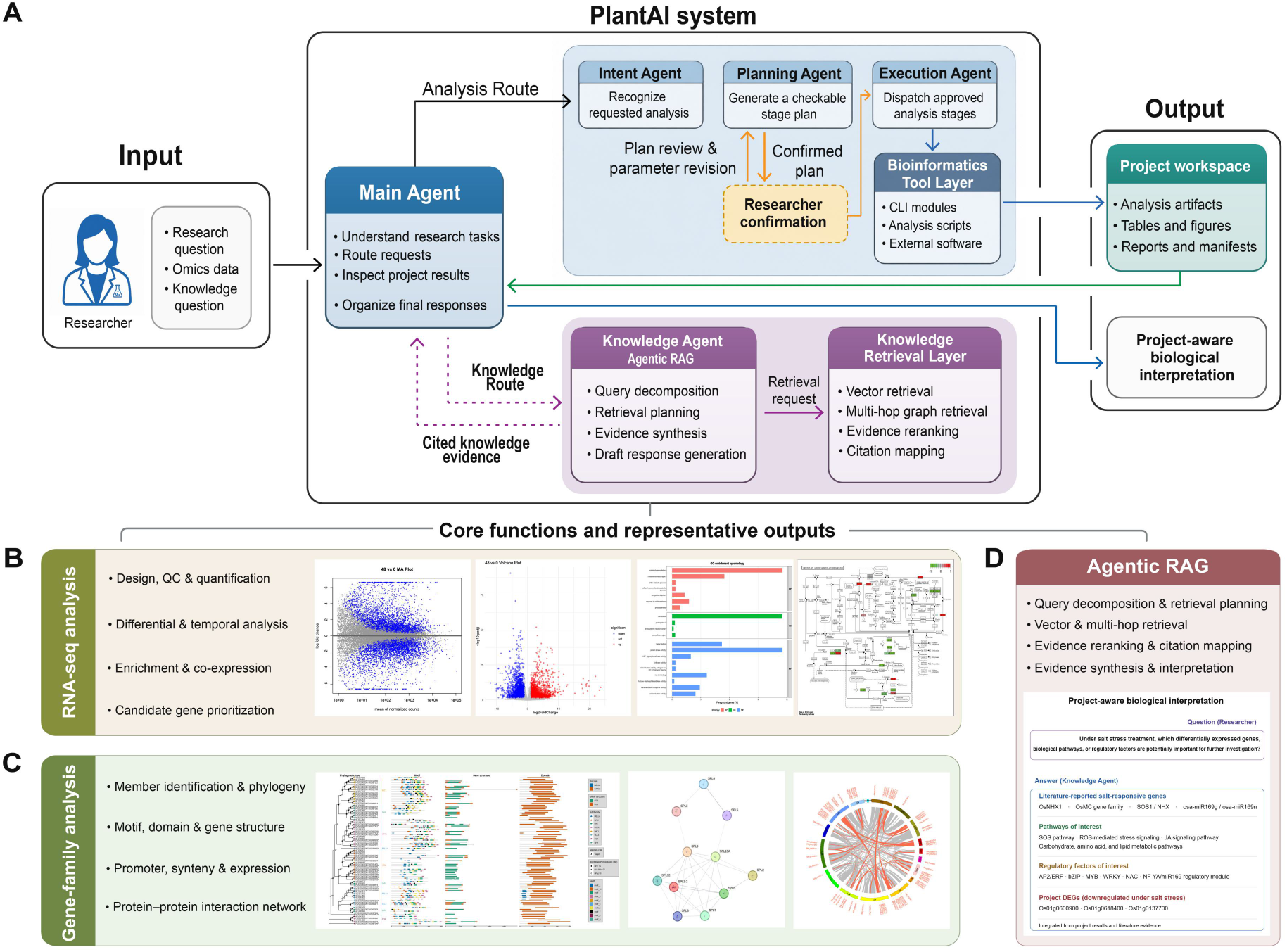
Overall architecture, core functions and representative outputs of PlantAI. A, The Main Agent coordinates the Analysis Route, Knowledge Route and Project workspace. B, RNA-seq analysis and representative outputs. C, Gene-family analysis and representative outputs. D, Knowledge Q&A and project-aware interpretation supported by PlantAI-RAG.

All analysis files, figures, reports and run records for a task are stored in an independent Project workspace and can be returned to the Main Agent for subsequent tasks or interpretation. PlantAI supports result traceability through stage-level records: run manifests record the analysis stage, execution status, parameter summary, key metrics and artifact paths; command and environment records document how the task was invoked and in which software environment; execution logs preserve the process, standard output, error messages and failure locations for each step; and intermediate and final artifacts retain the data underlying the analysis. By recording ‘what the result is’ separately from ‘how the result was generated’, these materials allow researchers to inspect, reuse and rerun the same analysis.

### 2.2 PlantAI-RAG and evidence-grounded knowledge interpretation

PlantAI-RAG is the plant-science specialist agent in PlantAI’s Knowledge Route and supports both general knowledge Q&A and supplementary interpretation of project results. Its knowledge base currently contains 31,207 plant-science literature records. After structural parsing, entity and relational-assertion extraction, entity normalization and evidence validation, these records yielded approximately 3.82 million canonical entities and 8.25 million literature-supported relational assertions, which were written to the Neo4j knowledge graph and Qdrant vector index, respectively (Fig. 2A). For project-aware Q&A, the Main Agent first reads analysis results from the Project workspace and then invokes PlantAI-RAG to retrieve relevant literature. The final response distinguishes among project-derived results, traceable external knowledge and interpretations jointly supported by both, thereby avoiding the presentation of literature knowledge as a new project finding or of a computational association as an experimentally validated causal mechanism.

**Figure 2.**
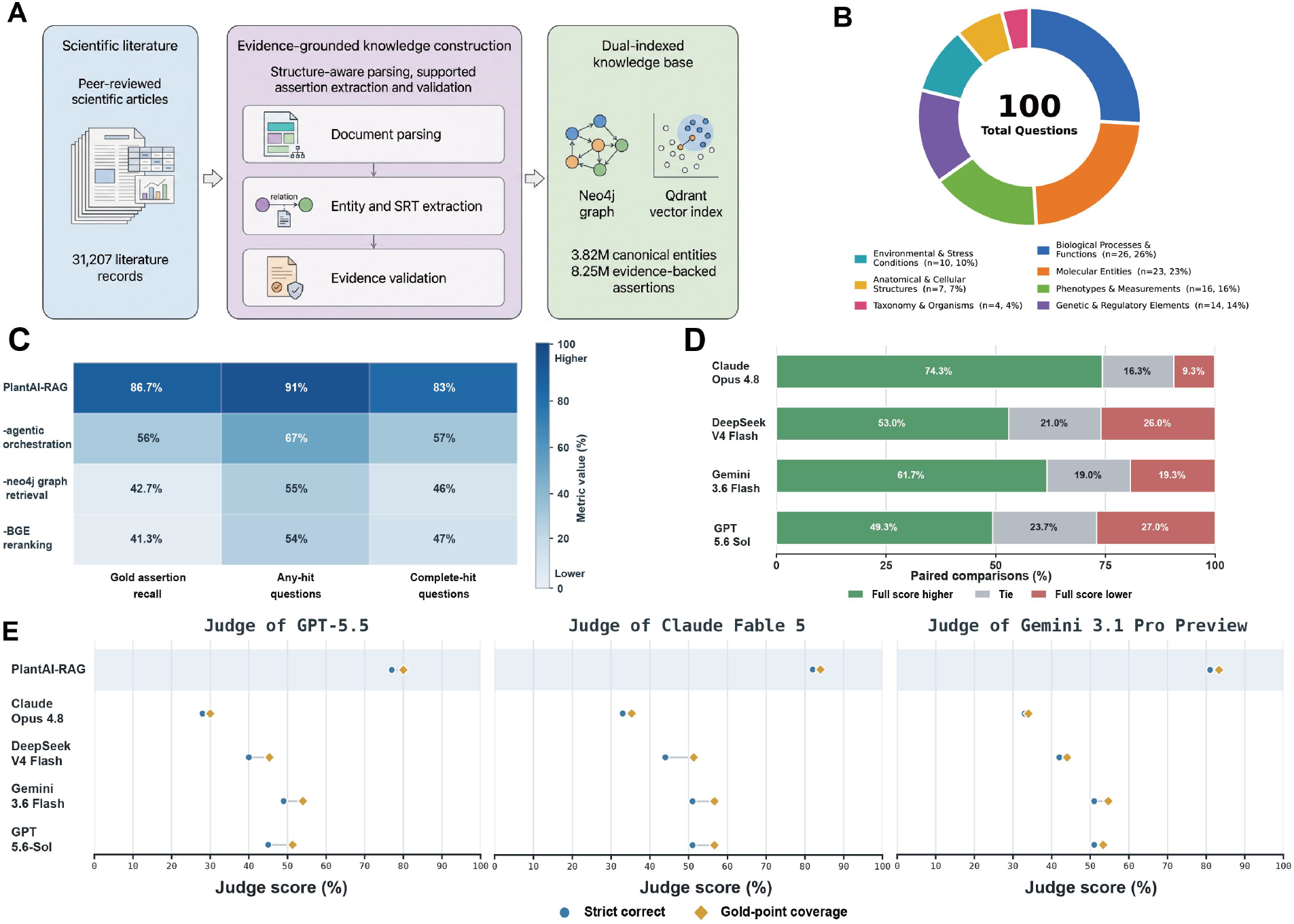
Plant-science evidence knowledge base and retrieval-augmented question-answering evaluation for PlantAI-RAG. A, Construction of the plant-science evidence knowledge base. B, Plant-science topic composition of the 100 evaluation questions. C, Ablation analysis of PlantAI-RAG. D, Question-level pairwise comparison of PlantAI-RAG with four general-purpose large language models. E, Strict correctness and Gold answer-point coverage assigned to each system by three independent judge models.

We evaluated PlantAI-RAG using 100 questions spanning seven plant-science topics (Fig. 2B). Its Gold evidence-assertion recall, Gold evidence hit rate and complete Gold evidence coverage rate were 86.7%, 91% and 83%, respectively. Removing agentic orchestration reduced these metrics to 56%, 67% and 57%; removing Neo4j graph retrieval reduced them to 42.7%, 55% and 46%; and removing BGE reranking yielded values of 41.3%, 54% and 47% (Fig. 2C). These changes indicate that question decomposition and retrieval planning, graph-relation retrieval and evidence reranking all contribute to the retrieval of necessary evidence. In question-level pairwise comparisons with four general-purpose large language models that did not use this knowledge-retrieval workflow, PlantAI-RAG had a higher win rate than loss rate against every model (Fig. 2D). Three independent judge models assigned strict correctness rates of 77%–82% and Gold answer-point coverage of 80%–84% (Fig. 2E). These results characterize the evidence-retrieval and answer performance of PlantAI-RAG on the current evaluation set and do not represent general accuracy across all plant-science questions.

### 2.3 Rice case study from RNA-seq to HXK/HKL family analysis and knowledge interpretation

We used 24 publicly available RNA-seq libraries from salt-tolerant rice HH11 and salt-sensitive rice IR29 sampled at 0, 6, 24 and 48 h after treatment with 200 mM NaCl to demonstrate a cross-module task ^9^ (Fig. 3). PlantAI completed quality control, alignment, quantification, differential expression analysis, functional enrichment, co-expression analysis and candidate-family screening, achieving a mean alignment rate of 89.47%. Carbohydrate metabolism emerged across multiple analysis outputs, OsHXK8 mapped to the hexokinase node in the KEGG starch and sucrose metabolism pathway, and its expression was consistently upregulated at all three post-treatment time points.

**Figure 3.**
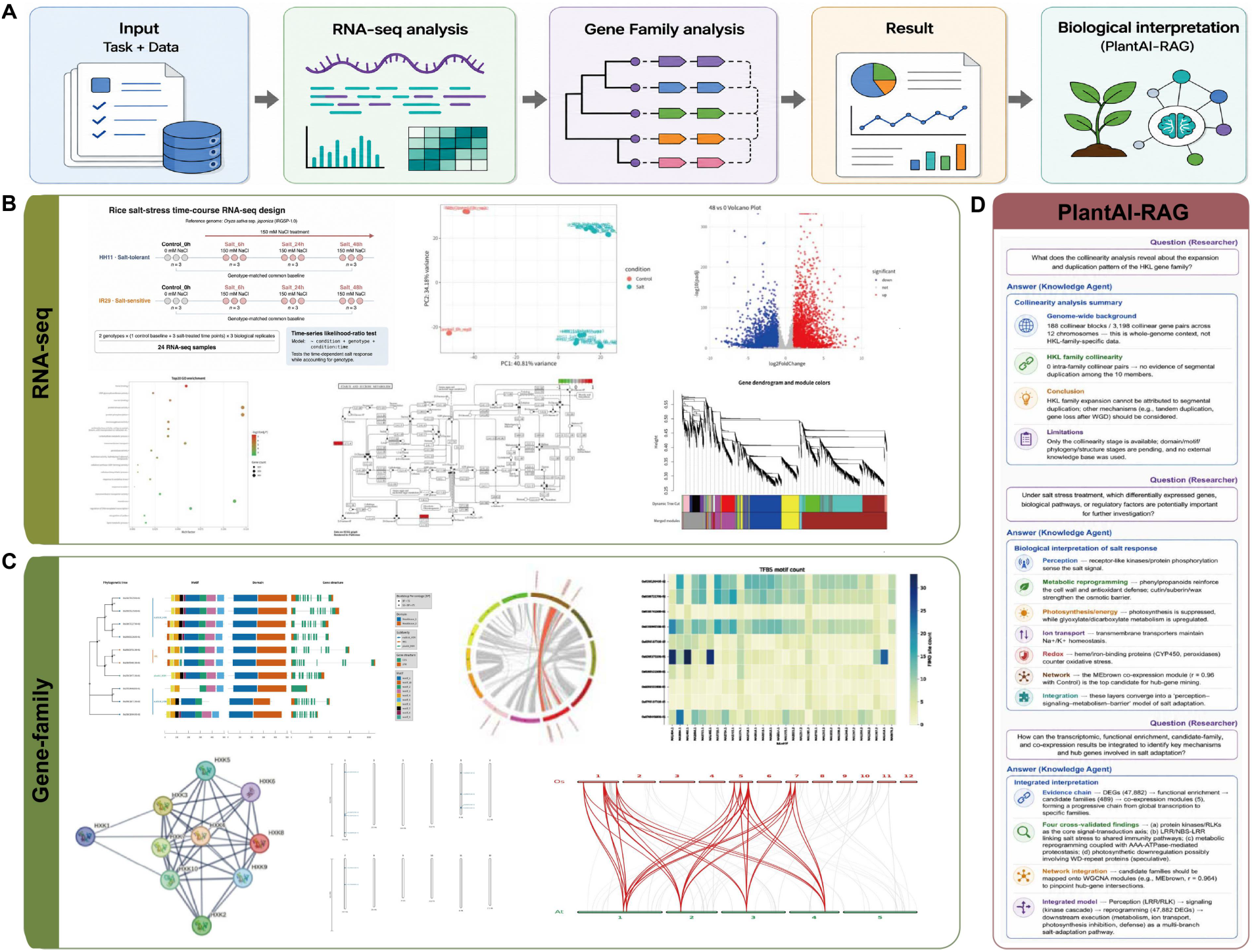
End-to-end rice salt-stress application in PlantAI. A, Experimental design and analysis workflow of the main case, showing how time-course RNA-seq data for HH11 and IR29 under salt stress proceed to candidate-family screening, HXK/HKL gene-family analysis and project-specific biological Q&A. B, Representative RNA-seq results, including sample principal component analysis, a differential-expression volcano plot, GO enrichment, KEGG pathways and WGCNA co-expression modules. C, Representative HXK/HKL gene-family results, including the integrated phylogeny–motif–domain– gene-structure plot, synteny, transcription factor-binding sites (TFBSs) and a predicted protein functional association network. D, Biological interpretation generated by PlantAI-RAG using the current project’s differential-expression, pathway and gene-family analysis results.

PlantAI subsequently identified HXK/HKL members containing both PF00349 and PF03727 core domains from the rice proteome and generated results for phylogeny, conserved motifs, gene structure, domain architecture, synteny, promoters and a predicted protein functional association network. The Main Agent read the run manifests, key tables and figures from each stage; organized the expression changes, pathway location and family-level results for OsHXK8 as project context; and then called PlantAI-RAG to retrieve literature evidence on sugar metabolism, sugar signalling and salt-stress responses. This case shows that PlantAI can connect RNA-seq, candidate-family screening, gene-family analysis and knowledge retrieval as a continuous task. The evidence supports prioritizing OsHXK8 for further validation but does not demonstrate that OsHXK8 determines salt tolerance in rice.

## 3. Discussion

PlantAI is intended not to reinvent algorithms for RNA-seq or gene-family analysis, but to organize research-task understanding, analysis planning, tool invocation, artifact management and knowledge retrieval within a single multi-agent system. Similar to systems such as BOLE^7^, PlantAI addresses the organization and traceability of established analytical methods in practical use. Its distinguishing feature is the inclusion of PlantAI-RAG for plant science, which allows actual project results to re-enter subsequent Q&A and literature-based interpretation. Run manifests, execution logs, command and environment records, and intermediate artifacts together provide an inspectable execution trail, reducing separation between analytical results and the processes that generated them.

The evidential boundaries of this study are also explicit. The rice case demonstrates an operational cross-module research chain; it does not validate a new salt-tolerance mechanism or establish that PlantAI outperforms other systems. The current demonstration is limited to one public dataset, lacks time-matched untreated controls after 0 h, and uses candidate-family screening that depends on available annotations and weighting choices. Promoter, TFBS and functional association network results are also primarily computational predictions. RAG responses are further affected by literature coverage, entity normalization, retrieval version and gene-identifier mapping. PlantAI should therefore be viewed as a computational workspace that helps researchers build traceable and reusable analysis chains that can be readily subjected to further validation. Key biological conclusions must still be evaluated against the underlying results, literature sources and experimental validation.

The range of analyses currently supported by PlantAI remains limited, focusing primarily on the RNA-seq, gene-family analysis and knowledge interpretation presented here. Future work will progressively integrate additional types of plant bioinformatics analysis and improve data handoffs, run records and result reuse across analysis stages. In parallel, PlantAI-RAG will continue to expand its coverage of plant and related biological literature, update evidence-backed assertions and entity relationships, and strengthen the literature-evidence and knowledge foundation for plant functional genomics, thereby supporting a broader range of project-specific questions and biological interpretations.

## 4. Conclusion

We present PlantAI, a multi-agent system for plant functional genomics. PlantAI uses the Main Agent to connect analysis planning, bioinformatics tools, the Project workspace and PlantAI-RAG, while preserving an auditable execution trail through run manifests, command and environment records, execution logs and analytical artifacts. The rice case shows how the system organizes RNA-seq, candidate-family screening, HXK/HKL gene-family analysis and knowledge retrieval into a continuous task. PlantAI provides an accessible, traceable and extensible way of working in plant bioinformatics.

## 5. Methods overview

### 5.1 System execution and run records

The Main Agent routes user requests to either the Analysis Route or the Knowledge Route. Analysis tasks proceed through intent recognition, plan generation, researcher confirmation and stage execution, and project files are organized according to stage-specific input-output contracts. Each completed stage generates a structured run manifest; associated tasks also preserve command scripts, software-environment information, execution logs and verifiable artifacts. Failed tasks display their status and a concise explanation in the frontend, while detailed records remain in the logs. Run manifests identify stage status, parameter summaries, result metrics and artifacts, whereas logs reconstruct the detailed execution process; the two record types are complementary rather than interchangeable.

### 5.2 PlantAI-RAG and the knowledge base

Plant-science literature is structurally parsed, followed by entity recognition, relational-assertion extraction, entity normalization and evidence validation, before being written to Neo4j and Qdrant. Knowledge Q&A begins with entity recognition, question decomposition and retrieval planning; vector retrieval, multi-hop graph expansion, evidence reranking and citation mapping are then combined to construct the answer context. Project-aware interpretation additionally receives result summaries extracted by the Main Agent from the Project workspace. Project files and run manifests remain the authoritative sources for project statistics; PlantAI-RAG supplies only external knowledge and source support.

### 5.3 RNA-seq analysis and candidate-family screening

The RNA-seq data were obtained from the public study by Fang et al. ^9^ and analysed using the Oryza sativa IRGSP-1.0 reference genome. Raw paired-end FASTQ data were first processed with fastp for adapter detection, quality trimming and length filtering, and quality-control statistics were then summarized. The resulting reads were aligned to the reference genome with HISAT2 ^10^; featureCounts generated gene-level raw counts ^11^, and Salmon quantified transcript abundance ^12^. Quantification results were also used for principal component analysis and checks of sample distances and correlations. DESeq2 performed differential expression analysis from raw counts^13^, while GO, KEGG, rank-based gene set enrichment analysis and WGCNA were used to organize functional and network evidence ^14–17^. Candidate families were ranked for review using evidence from differential responses, family response proportions, concordance of expression direction, co-expression and functional enrichment. Researchers made the final choice of study target after considering family size and analytical tractability.

### 5.4 HXK/HKL family analysis

The HXK/HKL family was defined by the joint presence of PF00349 and PF03727 ^18^. HMMER hmmsearch was first used to search the rice proteome with the two Pfam profile HMMs ^19^, while BLAST+ performed homology searches using reference family proteins ^20^. After candidate sets were combined, the dual-domain composition was verified with InterProScan/Pfam to determine family membership ^18,21^. MAFFT and IQ-TREE were used for multiple-sequence alignment and phylogenetic inference, respectively ^22,23^, and MEME was used for conserved-motif analysis ^24^. Pfam and GFF3 were used to organize domain and gene-structure information, MCScanX-based workflows were used for synteny analysis ^25^, and the 2-kb upstream regions were used for PlantCARE cis-regulatory element annotation and JASPAR Plants TFBS prediction ^26^. STRING was used to construct a predicted protein functional association network. All predictions were treated as candidate evidence rather than direct evidence of physical interactions or causal functions.

## Data availability

Raw rice RNA-seq data are available through NGDC BioProject PRJCA017851 and GSA CRA011547, with sequencing runs CRR809753–CRR809776.

## Acknowledgements

This work was supported by the Biological Breeding—National Science and Technology Major Project (2023ZD04073).

## Author contributions

Zhiqiang Xia and Tingxun Wu jointly conceived the study and designed the research programme (Conceptualization). Tingxun Wu developed the overall multi-agent system architecture and implemented the core algorithms (Software and Methodology). Zhuang Yang was responsible for debugging, functional validation and performance optimization of the PlantAI-RAG module (Validation and Software). Tingxun Wu wrote the initial manuscript draft; Zhiqiang Xia and Zhuang Yang systematically reviewed and revised the manuscript and contributed additional scholarly content. All authors read and approved the final manuscript.

